# A scalable approach to absolute quantitation in metabolomics

**DOI:** 10.1101/2024.09.09.609906

**Authors:** Luke S. Ferro, Alan Y. L. Wong, Jack Howland, Ana S. H. Costa, Jefferson G. Pruyne, Devesh Shah, Joshua D. Lauterbach, Steven B. Hooper, Mimoun Cadosch Delmar, Jack Geremia, Timothy Kassis, Naama Kanarek, Jennifer M. Campbell

## Abstract

Mass spectrometry-based metabolomics allows for the quantitation of metabolite levels in diverse biological samples. The traditional method of converting peak areas to absolute concentrations involves the use of matched heavy isotopologues. However, this approach is laborious and limited to a small number of metabolites. We addressed these limitations by developing Pyxis^TM^, a machine learning-based technology which converts raw mass spectrometry data to absolute concentration measurements without the need for per-analyte standards. Here, we demonstrate Pyxis performance by quantifying metabolome concentration dynamics in murine blood plasma. Pyxis performed equivalently to traditional quantitation workflows used by research institutions, with a fraction of the time needed for analysis. We show that absolute quantitation by Pyxis can be expanded to include concentrations for additional metabolites, without the need to acquire new data. Furthermore, Pyxis allows for absolute quantitation as part of an untargeted metabolomics workflow. By removing the bottleneck of per-analyte standards, Pyxis allows for absolute quantitation in metabolomics that is scalable to large numbers of metabolites. The ability of Pyxis to make concentration-based measurements across the metabolome has the potential to deepen our understanding of diverse metabolic perturbations.

## Introduction

The human body contains a vast and largely uncharacterized universe of small molecules^1^. Recent estimates suggest that the number of these small molecules, known as metabolites, in the body exceeds 200,000^2^. To begin to understand their functions in human biology, they must first be detected and quantified, a task undertaken by metabolomics. In liquid chromatography-mass spectrometry (LC-MS)-based metabolomics, metabolites are separated using chromatography and detected by a mass spectrometer, resulting in a characteristic peak for each metabolite. Quantitation of these peak areas provides information about metabolite abundance and can be either relative or absolute^3^. Relative quantitation involves comparing peak areas across different sample groups, without an explicit need to identify the molecule. However, peak areas cannot be directly compared without normalization^4,5^. Absolute quantitation involves converting peak areas into concentration units, such as µM or mols/10^6^ cells, using per-analyte standards. Concentration measurements provide a mechanism to reduce variability in large-scale studies and enable comparison between studies^6^. In addition, knowing metabolite concentrations is key to deciphering the roles they play in biological systems, such as whether a metabolite will bind to a cognate enzyme or receptor with known affinity^7^.

Despite the benefits of absolute over relative quantitation, the latter approach is far more common in metabolomics studies. This is because it is very difficult to obtain concentration values even for a small number of metabolites. The typical approach to absolute quantitation in metabolomics involves creating calibration curves for each analyte^8,9^. While this method is straightforward, creating large numbers of calibration curves is expensive, error-prone and laborious. In addition, external calibration curves do not account for matrix effects, which can significantly influence peak area measurements^10^. In mass spectrometry, the term ‘matrix’ refers to the biological sample type, such as cell lysate, containing the analytes. Matrix effects occur when co-eluting compounds from the matrix influence the detection of a target analyte, often leading to signal suppression between 60% and 90%^11^. Matched heavy isotopologues can be used to correct for matrix effects during concentration determination; however, they are expensive and available for only a small set of metabolites^12^. Thus, the traditional approach to absolute quantitation is difficult or impossible to scale to large metabolite numbers.

The need to utilize standards for each analyte also prevents the use of absolute quantitation in untargeted metabolomics studies. Unlike targeted studies, which investigate a pre-defined set of metabolites and utilize either relative or absolute quantitation, untargeted studies focus on discovering and annotating unknown compounds in the sample^3,13^. These hypothesis-generating experiments are essential for gaining mechanistic insight into perturbations that affect metabolism as well as for biomarker discovery, which requires looking at a broad swath of the metabolome^14^. However, as metabolite identity is not known prior to the experiment in untargeted metabolomics, quantitation is always relative. This is a significant limitation, as metabolite concentration information is crucial for data interpretation and prioritizing follow-up experiments in untargeted metabolomics.

Given the current challenges in metabolite quantitation, there is a pressing need for an approach to absolute quantitation in metabolomics that eliminates the need for per-analyte standards. Machine learning has emerged as a useful tool in diverse areas of biological data analysis^15^. However, most applications of machine learning in metabolomics have focused on annotating molecular identity rather than concentration prediction^16–22^. Ideally, a workflow would be able to both identify and provide concentration measurements for every detected metabolite. To address this gap, Matterworks developed Pyxis, a machine learning-based approach to absolute quantitation in metabolomics which eliminates the need for per-analyte standards and heavy isotopologues. Pyxis integrates an optimized LC-MS method for data collection, a machine learning model for concentration prediction and a cloud-deployed software for model inference.

The Pyxis model was built using a transformer backbone^23^ and trained on a dataset of labeled concentration-analyte pairs, with metabolites drawn from a large diversity of metabolic pathways. This dataset was collected in-house and, to our knowledge, represents the largest of its kind. A key feature of Pyxis is the use of StandardCandles^TM^, a set of internal standards utilized during training and inference which broadly function as universal standards. The StandardCandles enable the model to make concentration predictions across different instruments and instrument models. While proprietary, the StandardCandles are provided as part of a kit so that the platform can be set up in any laboratory with a high-resolution mass spectrometer. In addition, we developed a web-based software to support running large numbers of samples through the model.

In this study, we demonstrate Pyxis performance using a mouse blood plasma dataset collected at Boston Children’s Hospital (BCH). Pyxis successfully quantified the concentration dynamics of over 100 plasma metabolites, demonstrating its ability to capture biological changes in animal models of disease. Pyxis results were validated through comparison to both orthogonal concentration measurements as well as an independent dataset of the same samples analyzed using relative quantitation. When used for targeted quantitation, we demonstrate that Pyxis can report concentrations for additional metabolites without the need to collect additional sample data. This is in contrast to traditional methods of absolute quantitation. We also show that Pyxis can provide absolute quantitation as part of an untargeted analysis workflow, fundamentally changing the type of output possible in untargeted metabolomics. The ability of Pyxis to report metabolite concentration without the need for per-analyte standards can allow for the scope of absolute metabolomics studies to be greatly expanded. Together, these data demonstrate how machine learning can enable scalable absolute quantitation for metabolomics in a broad range of metabolomics studies.

## Results

### Pyxis provides insight into metabolome concentration dynamics in blood plasma

Metabolomic analysis was performed on samples from two cohorts of mice fed diets with different levels of folate, a vitamin that plays an essential role in one-carbon metabolism^24^. Mice were divided into four distinct groups, delineated by sex and diet (**Fig. 1a**). Blood was collected via cheek bleed and processed at weekly time points up to 6 weeks. **Supplementary table 1** details the number of mice in each cohort and the volume harvested at each time point. Overall, there were 7 male and 7 female mice, with 3-4 mice in each diet cohort.

**Fig. 1.**
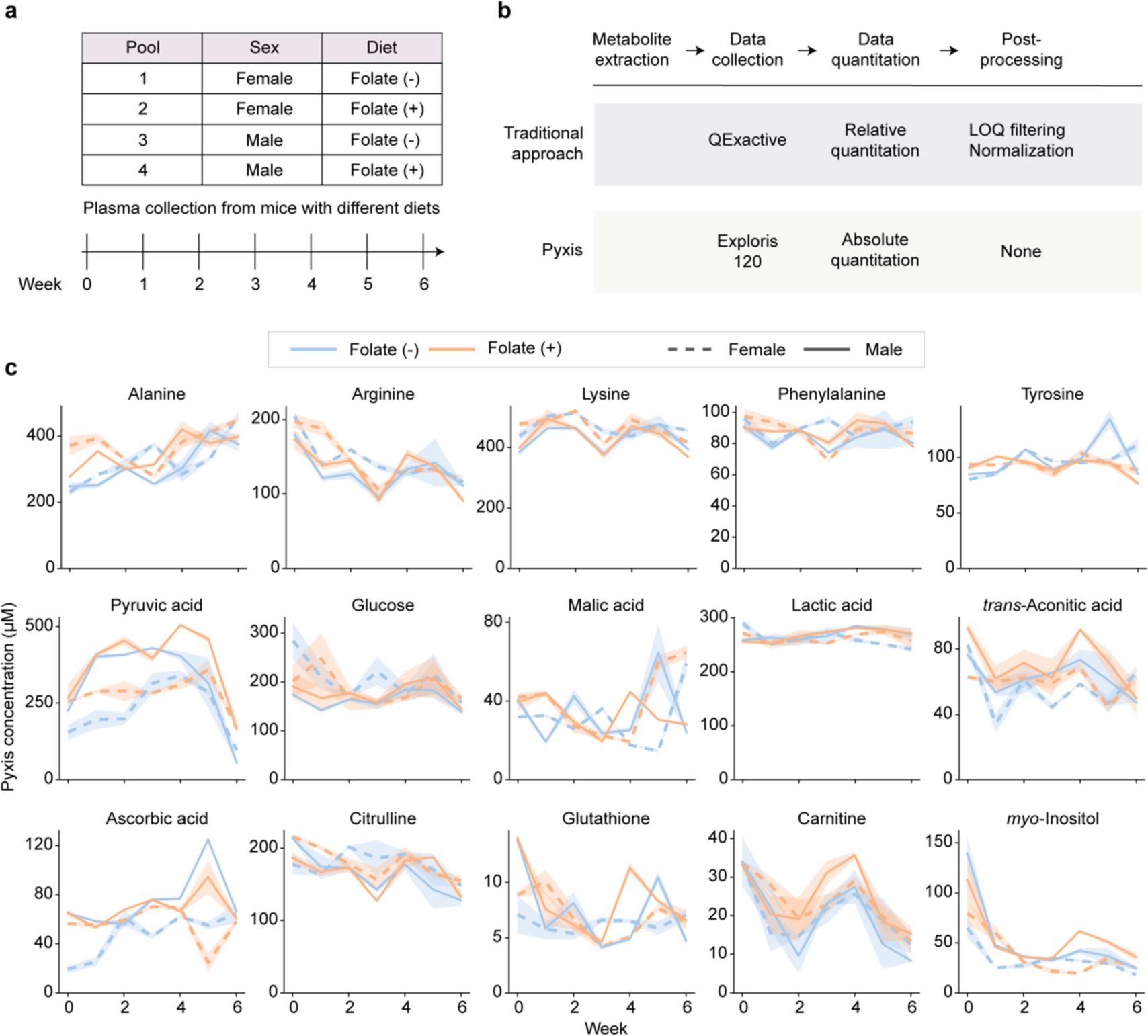
Pyxis provides insight into metabolome concentration dynamics in murine plasma. **a** Schematic of the experimental groups. Blood was harvested from mice of different sex and diets over the course of 6 weeks and plasma was immediately isolated. Plasma samples were combined into 4 distinct pools. **b** Data were collected and analyzed with two different approaches. Each approach differed in the methods underlying data collection, quantitation, and processing. **c** Pyxis concentration data for a subset of quantified metabolites. Orange and blue lines depict the folate-replete and folate-deficient diets, respectively. Solid and dotted lines differentiate the male and female mice, respectively. The shaded area depicts standard deviation between 2 technical replicates. Concentration values are adjusted for the dilution factor used during sample preparation.

After sample collection had finished, the samples were split and analyzed via independent workflows involving either relative or absolute quantitation (**Fig. 1b**). Relative quantitation was carried out using a traditional targeted LC-MS-based approach, which involved methanol-based plasma extraction, pHILIC chromatography, and analysis with an Orbitrap mass spectrometer. For absolute quantitation, Pyxis model v0.10.0 was used which provides quantitation of 159 metabolites, along with a proprietary quality control (QC) analyte. We conducted two types of QC for the absolute quantitation experiment: one for the raw data and another for model performance. QC of the raw data was performed using signals from the StandardCandles and a set of spiked-in heavy isotoplogue standards, while QC of model performance was accessed using the proprietary QC analyte (**Supplementary Fig. 1**).

Across the samples, 103 metabolites were detected and determined to be within the quantitative range of the model, allowing for absolute concentrations to be obtained (**Supplementary Data 1**). The metabolites quantified by Pyxis covered a wide range of metabolic pathways, including amino acid metabolism, glycolysis, and redox pathways (**Fig. 1c**). The Pyxis data revealed that amino acid concentrations in mouse plasma remained consistent over weeks during exposure to a folate-deficient diet and aligned well with previously published values (**Supplementary Table 2**)^25,26^. Other metabolites showed notable changes over the course of the experiment. Pyruvic acid declined across all the experimental groups from weeks 4 to 6, regardless of dietary folate concentration. In contrast, carnitine levels oscillated over time, peaking at weeks 0 and 4. Although clear differences were not observed between diets, sex-specific metabolic variation highlighted the importance of considering sex as a variable in metabolomics studies^27^. Together, these data demonstrate that Pyxis can effectively capture the concentration dynamics of diverse metabolites over time in murine plasma, providing insight into metabolic processes and their regulation.

### Comparison of Pyxis data to orthogonal measurements and across institutions

For this dataset, we validated the performance of the Pyxis model by conducting several comparative analyses. For a subset of endogenous metabolites in the plasma samples, concentrations were determined in an orthogonal manner using spiked-in, matched heavy isotopologues. Pyxis concentration measurements were then compared to these orthogonal measurements, and we observed strong correlations between the two methods (**Fig. 2a, Supplementary Fig. 2, Supplementary Data 2**). For metabolites lacking matched heavy isotopologues, we looked at how the peak area trends compared to the Pyxis concentration trends. Here, peak areas are the area under the curve from the extracted ion chromatogram for each metabolite. For metabolites with clear trends of changes in the peak areas between samples, we also observed similar trends in the Pyxis results, providing confidence in the Pyxis predictions for these metabolites (**Supplementary Fig. 3**).

**Fig. 2.**
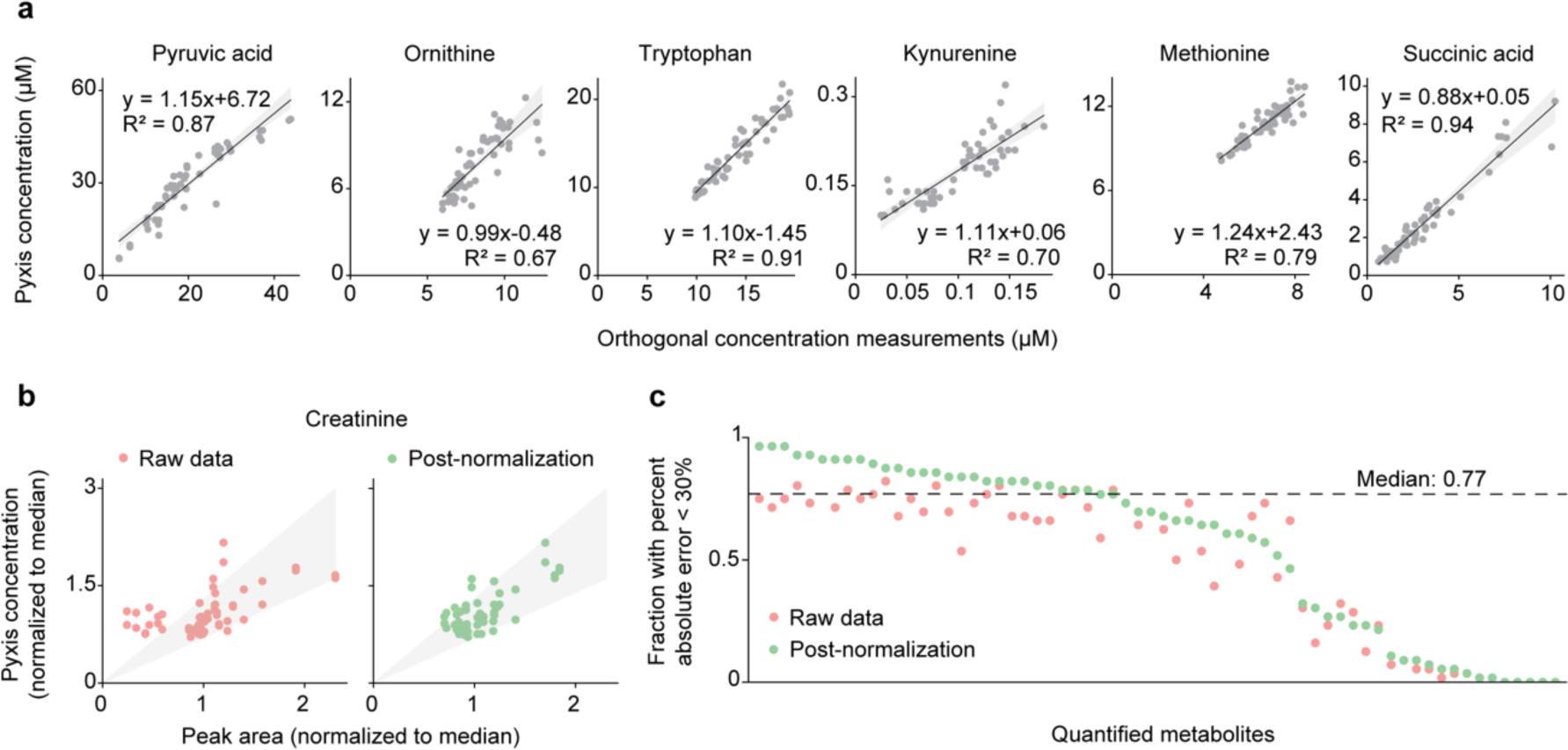
Comparing Pyxis data to orthogonal measurements and across institutions. **a** Comparison of Pyxis concentration data to orthogonal concentration measurements for a set of endogenous metabolites in the plasma samples. Orthogonal concentration measurements for 22 metabolites were obtained using matched heavy isotopologues spiked-into the extraction solution. Linear fit equations and R-squared values are printed on the plots. Data for the remaining metabolites shown in Supplementary Fig. 2. **b** Comparison of Pyxis concentration data to peak area data collected at BCH. Pyxis concentration and peak area data for each metabolite are normalized to their median value to allow for comparison. Raw peak area data from BCH are shown in red, normalized peak area data are given in green. Normalization given in the color legend refers to the post-acquisition data processing used to reduce injection-to-injection variability in the relative quantitation dataset (described in the Methods). Creatinine is shown as an example of how normalization of the relative quantitation data affects the correlation between the two quantitation methodologies. The shaded grey area represents +/− 30% band with respect to the normalized peak area data. **c** Data showing the fraction of the normalized Pyxis data for each metabolite that had a percent absolute error value less than 30%, with respect to the normalized relative quantitation data.

Additional data were collected in parallel at BCH and analyzed using traditional relative quantitation. These data were acquired using the same samples, but a different LC-MS system and method. Data processing involved several additional normalization steps to reduce injection-to-injection variability. Both raw and post-normalization data from the relative quantitation workflow were compared to the Pyxis data (**Fig. 2b-c**). As relative and absolute quantitation data cannot be directly compared, both sets of data were normalized with respect to their median values. The fraction of model predictions for each metabolite which had a percent absolute error less than 30% was determined. We found that the median percent absolute error value across all metabolites was 77% after normalization (**Fig. 2c**), meaning that 77% of the Pyxis data matched to within +/− 30% of the relative quantitation data. These results demonstrate both the need to normalize data in relative quantitation and the strong performance of Pyxis when compared to traditional methods of quantitation.

### Absolute quantitation on additional metabolites with previously confirmed identity

The relative quantitation dataset (**Fig. 2**) included 40 metabolites not included in the output of Pyxis model v0.10.0. We sought to expand the set of metabolites comparable across institutions by having Pyxis report the concentrations of these 40 metabolites. In the traditional approach to absolute quantitation, converting additional peak areas to concentrations would require purchasing standards and collecting calibration curve data for each additional metabolite. With Pyxis, additional concentration measurements can be obtained without the need to collect additional data.

We reprocessed the original raw data with Pyxis, after providing the model with the chemical formula and SMILES (Simplified Molecular Input Line Entry System) of the additional metabolites. SMILES are a standardized way to represent chemical structures as text^28^. Of the 40 additional metabolites, 29 were detected with the Pyxis LC-MS method and determined to be within the quantitative range of the model. Pyxis predictions were compared to peak area measurements from both Matterworks and BCH (**Fig. 3a-b, Supplementary Fig. 4**). We also confirmed identity using targeted MS/MS data collected with the Pyxis LC-MS method (**Fig. 3c-d, Supplementary Fig. 4**).

**Fig. 3.**
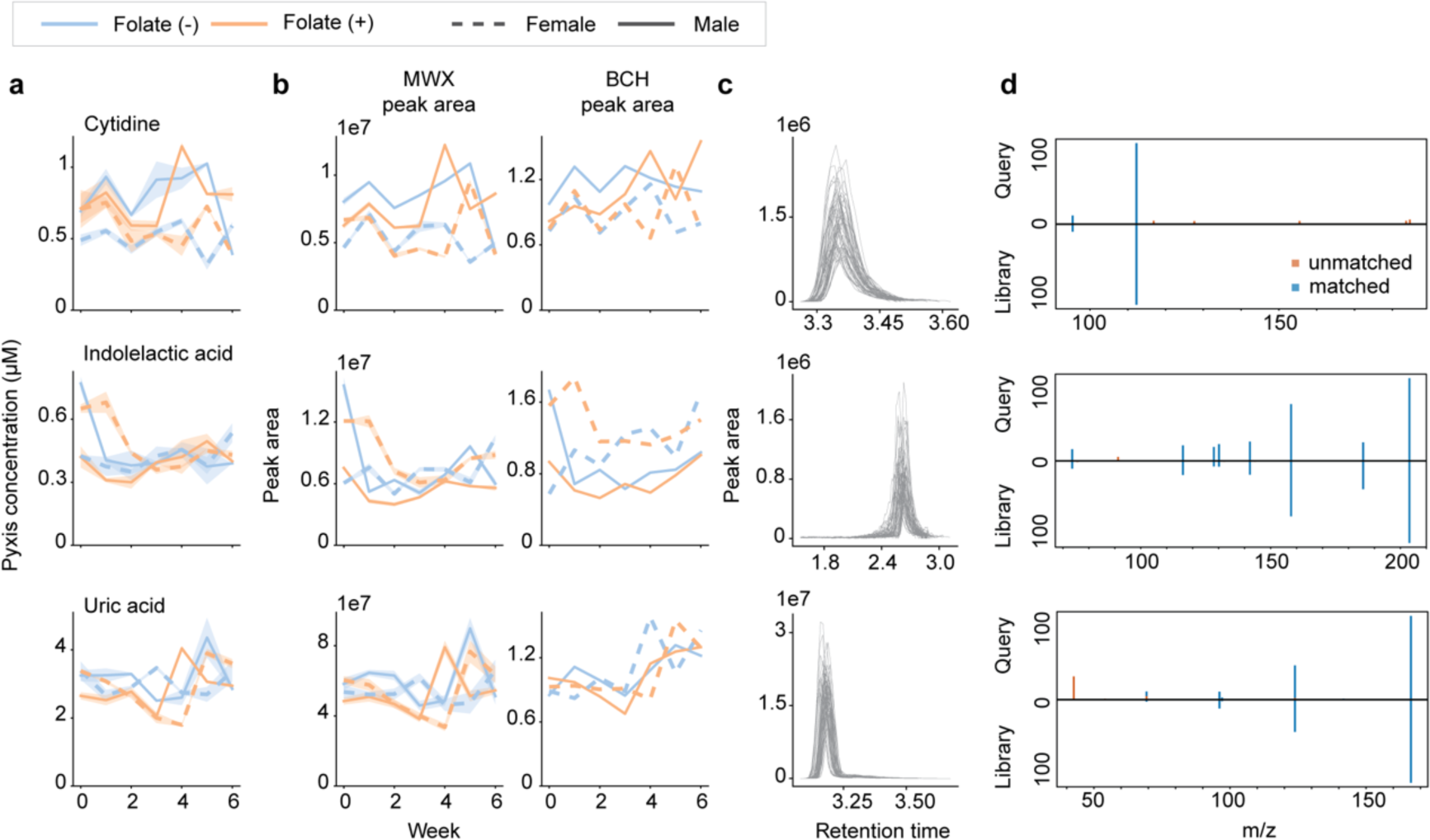
Absolute quantitation on additional metabolites with previously confirmed identity. **a** Pyxis concentration measurement for each metabolite. Legend given above the plots. **b** Peak area measurements (arbitrary units) given for Matterworks (MWX) and BCH. The shaded area depicts standard deviation between 2 technical replicates. **c** Peak area plots for each metabolite in the pooled sample. **d** MS/MS data mirror plots for each analyte. Top half is the query (acquired) spectrum, bottom half is the library spectrum. Blue bars are those fragments in which there was a library match. Orange bars are unmatched fragments. Data were acquired from pooled plasma samples.

To validate model performance on the 29 metabolites detected with the Pyxis LC-MS method, we analyzed a pooled plasma dilution series with Pyxis. In a dilution series, there should be a linear relationship between dilution factor and Pyxis concentration predictions. Thus, data was filtered based on linearity to reveal metabolites with high-confidence concentration predictions. In the plasma pool, 16/29 quantified metabolites passed the linearity filter described in the **Methods** (**Supplementary Fig. 5**). Of note, metabolites can fail this filter if their signal is below the limit of quantitation for a significant portion of the dilution series. Importantly, pool dilution results test the ability of Pyxis to predict differences in absolute concentration rather than probing accuracy directly. Together, these data demonstrate the ability of Pyxis to extend the scope of absolute quantitation studies without the need to collect additional per-analyte calibration curves.

### Expanding metabolite coverage with absolute untargeted metabolomics

There are a number of traditional approaches to untargeted metabolomics analysis, supported by software such as MZmine^29^, MS-DIAL^30^, and XCMS^31^ as well as commercial software developed by instrument manufacturers. These tools are well-established and allow users to discover unknown metabolites in their samples using MS/MS data. More recent techniques which utilize machine learning, such as SIRIUS^32^, MIST^18^ and MS2Mol^17^, allow for a higher percentage of features to be annotated with molecular identity in a given study^33^. However, despite their diversity, the final output of all these approaches remains relative quantitation. The lack of concentration information is a fundamental limitation in current untargeted metabolomics studies^34^. Thus, we investigated the ability of Pyxis to provide absolute quantitation as part of an untargeted metabolomics workflow.

In our absolute untargeted metabolomics workflow, metabolite identity was determined using MZmine and concentrations determined by Pyxis (**Fig. 4a**). This workflow leverages the strengths of both targeted and untargeted metabolomics, combining the comprehensive characterization of untargeted methods with the absolute quantitation offered by targeted approaches. Using MS/MS data, this workflow identified and provided concentration measurements for 137 metabolites, 83 of which had not been reported in the original Pyxis dataset (**Fig. 4b-c**). Importantly, Pyxis can provide absolute concentrations for metabolites identified by any untargeted metabolomics software, not just MZmine. Confidence in the concentration results was determined by comparison to a pooled plasma dilution series, which showed that 47/83 metabolites displayed expected linear behavior (**Fig. 4d**). Together, these data show that Pyxis can provide concentration data for any set of identified metabolites, enabling the integration of absolute quantitation into untargeted metabolomics.

**Fig. 4.**
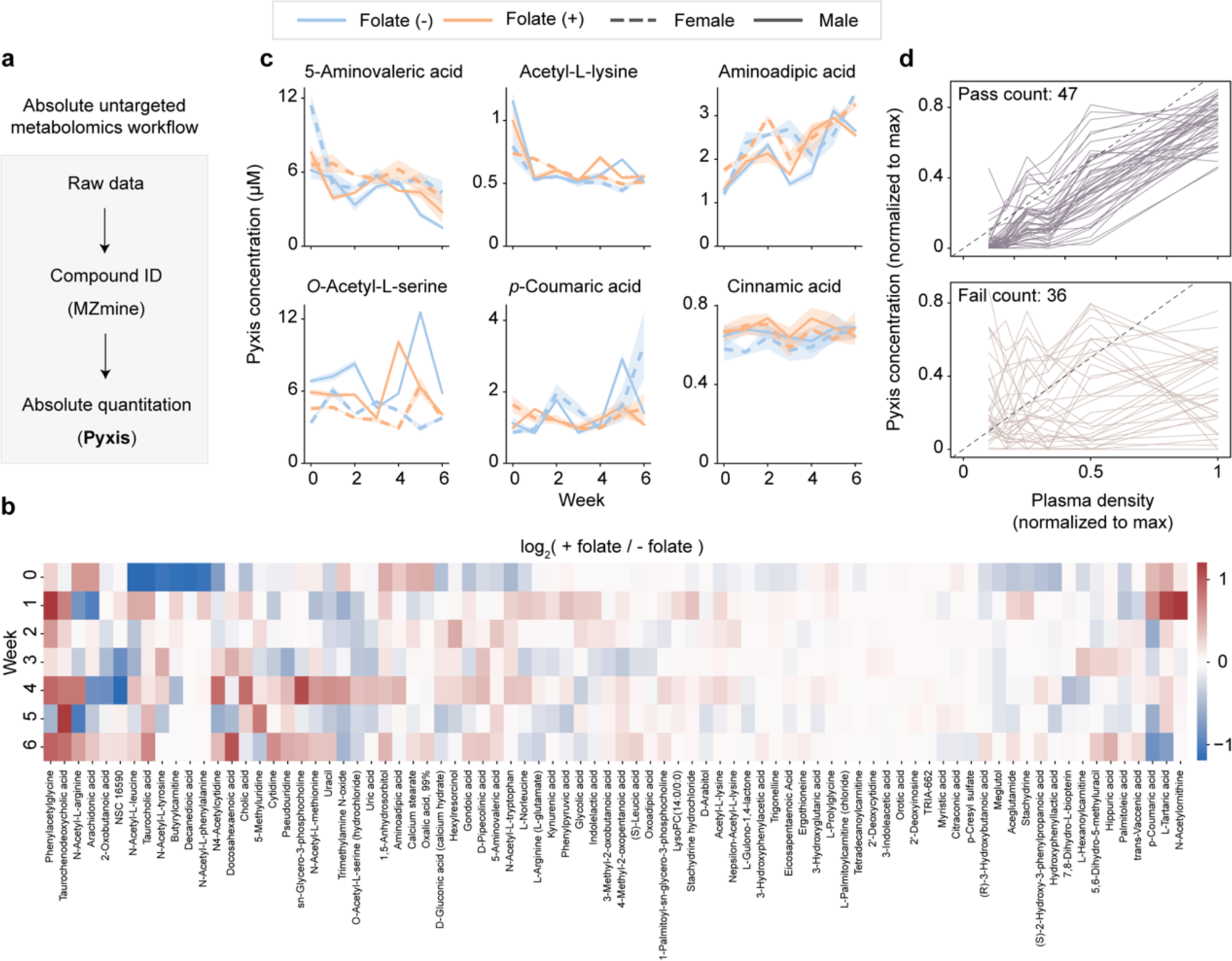
Expanding metabolite coverage with absolute untargeted metabolomics. **a** Diagram of the workflow for absolute untargeted metabolomics. Compound identity is established with MZmine, and absolute quantitation is performed by Pyxis. **b** Cluster map for the 83 metabolites identified in the absolute untargeted metabolomics workflow but not already captured in original data from Fig. 1. Depicted is the log_2_ fold change of the average Pyxis concentration for each metabolite for the (+) folate diet over the (-) folate diet over time. Legend given on the right of the plot. Each column represents a quantified metabolite over time in weeks, as given on the y-axis. **c** Pyxis quantitation of a subset of the metabolites shown in (b). Legend above the plots. The shaded area depicts standard deviation between 2 technical replicates. Subset chosen to depict metabolites with concentrations that increased, decreased or remained constant over time. **d** Pooled sample dilution series data for the 83 metabolites defined in (b). Metabolites that have passed or failed the linearity filter (described in Methods) given on the top and bottom plot, respectively.

## Discussion

In this work, Pyxis demonstrated the ability to accurately determine metabolite concentrations in mouse plasma, a matrix relevant to animal models of disease. Pyxis captured the dynamics of 103 metabolites, providing insight into a wide range of metabolic pathways. We validated model predictions using a traditional quantitation approach based on matched heavy isotopologues. In addition, comparison of data from an independent relative quantitation workflow to the Pyxis data further reinforced the accuracy of Pyxis data, with a significant portion of data matching within a 30% absolute error threshold. These results show that Pyxis provides a solution to the problems associated with absolute quantitation in metabolomics; namely, the creation of per-analyte calibration curves and the limited availability of heavy isotopologues. For targeted metabolomics studies, Pyxis enables reporting concentrations for all measured metabolites, regardless of the availability of standards.

Eliminating the need for matched standards allows for absolute quantitation to be utilized in untargeted metabolomics. Untargeted metabolomics studies yield a large number of hits, and it is often not possible to follow-up all of them. Unlike relative fold change data, which can be misleading if concentrations are very low, absolute quantitation provides critical insight into which hits may be the most biologically relevant. For microbiome studies, for example, a researcher may want to focus on secreted molecules that are within a concentration range that may influence human cell behaviors. In drug metabolism and pharmacokinetics (DMPK) or exposomics, the biological effects of drug breakdown products or environmental contaminants are also highly concentration dependent. Pyxis can be used as part of any untargeted metabolomics workflow to provide absolute quantitation of identified metabolites. Furthermore, reporting concentration values for untargeted metabolomics studies will enable data from different labs to be compared and combined, allowing for a clearer view of metabolism to be built over time.

Absolute quantitation with Pyxis also has the benefit of simplifying key aspects of metabolomics data processing. First, Pyxis removes the requirement for additional signal normalization. Adjusting for fluctuations in mass spectrometer response typically requires heavy isotopologue standards, pooled samples and custom data analysis pipelines^4^. Pyxis makes this process standardized and automated. Second, Pyxis automatically determines whether a metabolite signal is within the limit of quantitation. If metabolite levels are too low or high, a change in concentration will not be reflected in a change in instrument signal^3^. Pyxis is trained to only report metabolite concentrations that are within the limit of quantitation, without the need for expert intervention. Third, Pyxis accounts for matrix effects during concentration determination. Metabolite signals can be either suppressed or enhanced by the biological matrix^11^. Pyxis adjusts for these effects on a per-analyte level, allowing the researcher to move seamlessly from matrix to matrix.

The ability of Pyxis to scale to large numbers of metabolites will allow for absolute quantitation to be utilized more readily in areas such as personalized medicine and genome-scale modeling. In personalized medicine, metabolomics can help stratify patients using biomarkers^35^. However, robust biomarker discovery often requires data collected across different times and institutions to achieve the necessary statistical power^6^. Absolute quantitation provides a key mechanism to reduce variability in large scale studies but has previously been untenable for large numbers of metabolites. Similarly, genome-scale metabolic models (GEMs) can also benefit from absolute quantitation. GEMs capture metabolism at a broad scale and have proven useful in both medicine and biotechnology^36^.

Concentration measurements can be used in GEMs to relate metabolite levels to the binding affinities of their respective enzymes^7^. Pyxis addresses the need for high-throughput absolute quantitation in both of these domains. With Pyxis, data can be collected in less than seven minutes per sample and thousands of samples can be analyzed in minutes, as no peak picking or expert intervention is required.

Machine learning has already begun to reshape how biological research is performed, yet its potential remains to be realized in many domains. The power of a tool like Pyxis highlights the need to apply machine learning to new areas of biological data analysis. We show that Pyxis overcomes a long-standing challenge in the field of metabolomics, breaking the reliance on per-analyte standards for absolute quantitation. Looking forward, the continued integration of machine learning in metabolomics promises to not only enhance the scope and quality of metabolomics studies, but also help catalyze fundamental breakthroughs in our understanding of metabolism.

## Methods

### Animal work

The Institutional Animal Care and Use Committee (IACUC) at BCH approved all animal work carried out in this study (protocol #1727). At 5 weeks of age, NOD.CB17-*Prkdc*^scid^/NCrCrl (NOD-SCID) mice (originally purchased from Charles River Laboratories and maintained at the BCH mouse facility through in-house breeding) were assigned to either a folate-replete diet (TD.220361, Envigo) or a nutrient-matched folate-deficient diet (TD.220362, Envigo) ad libitum. Autoclaved water was provided ad libitum. Mice were weighed and monitored every 2-3 days for adverse effects of the diet, including: lethargy, drop in weight (10% over a day or 20% since the beginning of the experiment), lack of movement, hunched back, or hair loss.

### Sample harvest

Cheek bleeds were performed using 5 mm lancets (Fisher #NC9891620). Approximately 50 μL of mouse blood was collected in a heparinized collection tube (VWR 103093-113). Whole blood was spun at 2000 RCF for 10 mins at 4 °C to separate the plasma from red blood cells. The supernatant plasma was collected and frozen at –80 °C until use. Samples were thawed at the end of the experiment, pooled, and then divided into two aliquots for metabolomics analyses.

### Sample extraction – traditional relative quantitation

Pooled plasma (5 μL) was extracted using 200 μL of extraction buffer (80% LC-MS-grade methanol, 20% LC-MS water) and supplemented with isotopically labelled internal standards (17 amino acids and isotopically labelled reduced glutathione, Cambridge Isotope Laboratories, MSK-A2-1.2 and CNLM-6245-10), as well as aminopterin. After adding extraction buffer, samples were vortexed briefly (10 sec). Samples were then centrifuged for 10 min, 4 °C, at maximum speed on a benchtop centrifuge (Eppendorf) and the cleared supernatant was transferred to a new tube. Samples were dried using a nitrogen dryer while on ice, and special attention was given to minimize the time of drying and to not let samples idle in the dryer (Reacti-Vap™ Evaporator, Thermo Fisher Scientific, TS-18826) once the drying process was completed. Needles were continuously adjusted to the surface of the liquid as the samples evaporated to expedite the drying process. Dried samples were stored at –80 °C until able to be analyzed. Samples were then reconstituted in 20 µL LC-MS-grade water supplemented with QReSS (Cambridge Isotope Laboratories, MSK-QRESS-KIT) by brief 10s vortexing. Reconstituted samples were spun for 10 min at maximum speed on a benchtop centrifuge at 4 °C and the supernatant was transferred to LC-MS micro vials (National Scientific, C5000-45B). A small amount of each sample was pooled and serially diluted 3– and 10-fold to be used as quality controls throughout the run of each batch.

### Sample extraction – Pyxis

Plasma (10 μL) was extracted with 40 μL extraction solution (50%, 30% and 20% methanol, acetonitrile and water, respectively). Samples were vortexed at 2000 RPM for 10 minutes at 4°C and then centrifuged at 20,000 RCF for 10 minutes at 4°C. 45 μL of the supernatant was then transferred to a new tube and combined with 45 μL extraction solution containing a 2x working concentration of StandardCandles. After agitating the tube, samples were transferred to 96-well plates and the plates sealed.

The final concentration of StandardCandles in the sample was 5 μM. A proprietary QC analyte is included as part of the StandardCandles mixture, with a final concentration of 1 μM. A set of 22 matched heavy isotopologues were also included in the extraction buffer at a final concentration of 5 μM. The metabolites with matched heavy isotopologues were as follows: 2-oxopropanoic acid, citric acid, tryptophan, fumaric acid, glucose-6-phosphate, glycine, alanine, arginine, asparagine, aspartate, glutamate, glutamine, leucine, lysine, kynurenine, methionine, phenylalanine, proline, serine, threonine, valine, succinic acid. Heavy isotopologues are not utilized by Pyxis during model inference and were only included in this experiment to enable orthogonal concentration determination.

### LC/MS method – traditional relative quantitation

Reconstituted sample (2 µL) was injected into a ZIC-pHILIC 150 × 2.1 mm (5 µm particle size) column (EMD Millipore) operated on a Vanquish™ Flex UHPLC System (Thermo Fisher Scientific, San Jose, CA). Chromatographic separation was achieved using the following conditions: buffer A was acetonitrile; buffer B was 20 mM ammonium carbonate, 0.1% ammonium hydroxide. Gradient conditions were as follows: linear gradient from 20 to 80% B; 20–20.5 min: from 80 to 20% B; 20.5–28 min: hold at 20% B. The column oven and autosampler tray were held at 25 °C and 4 °C, respectively.

MS data acquisition was performed using a QExactive benchtop orbitrap mass spectrometer equipped with an Ion Max source and a HESI II probe (Thermo Fisher Scientific, San Jose, CA) and was performed in positive and negative ionization mode in a range of m/z = 70–1000, with the resolution set at 70,000, the AGC target at 1 × 10^6^, and the maximum injection time at 20 ms. For polar metabolites, HESI conditions were as follows: Sheath gas frow rate: 35; Aug gas flow rate: 8; Sweep gas flow rate: 1; Spray voltage: 3.5 kV (pos), 2.8 kV (neg); Capillary temperature: 320 °C; S-lens RF: 50; Aux gas heater temperature: 350 °C. For T3/T4 HESI conditions were as follows: Sheath gas frow rate: 40; Aug gas flow rate: 10; Sweep gas flow rate: 0; Spray voltage: 3.5 kV (pos), 2.8 kV (neg); Capillary temperature: 380 °C; S-lens RF: 60; Aux gas heater temperature: 420 °C.

### LC/MS method – Pyxis

Sample (4 µL) was injected into an Atlantis Premier BEH Z-HILIC VanGuard FIT Column (2.1 mm x 50 mm, 2.5 µm) coupled to a guard cartridge. Mobile phase A was 20 mM ammonium carbonate in water with 0.25% (v/v) ammonium hydroxide (pH 9.55). Mobile phase B was acetonitrile. The autosampler wash solutions were Wash 1 (aqueous) contains 95% water, 5% acetonitrile, and 0.1% formic acid; Wash 2 (organic) comprises 45% acetonitrile, 45% isopropanol, and 10% acetone. Additionally, a rear seal wash solution was used, consisting of 90% water and 10% methanol. All reagents were LC/MS grade.

The LC method was as follows: 0-1 min, 95% B; 1-8.5 min, ramp from 95% to 20% B; 8.5-9.5 min, 20% B; 9.5-10 min, ramp from 20% to 95% B. The flow rate was 0.5 mL/min. The autosampler was kept at 4 °C, and the analytical columns were held at room temperature. Data was collected using a Thermo Fisher Orbitrap Exploris 120 with settings as follows: probe positions (front-to-back position 2, rotational position at center, and spray insert depth M), static spray voltage (positive ion 3500 V, negative ion 2500 V), sheath gas 50, auxiliary gas 10, sweep gas 1, ion transfer tube 315°C and vaporizer temperature 350°C. The full scan data is collected from 0 to 6.7 minutes in polarity switching mode: RF lens 55%, maximum injection time 60 ms, scan range 70 to 800 m/z, resolution 60,000. Internal mass calibration with EASY-IC was used.

### Data analysis – traditional relative quantitation

Relative quantitation of polar metabolites was performed with TraceFinder 5.1 (Thermo Fisher Scientific, Waltham, MA) using a 5 ppm mass tolerance and referencing an in-house library of chemical standards. Pooled samples (1x) and fractional dilutions (0.33x, 0.1x) were prepared as quality controls and only those metabolites were taken for further analysis, for which the correlation between the dilution factor and the peak area was >0.95 (high-confidence metabolites) and for which the coefficient of variation was below 30%. Normalization for biological material amounts was based on the total integrated peak area values of high-confidence metabolites within an experimental batch after normalizing to the averaged factor from all mean-centered chromatographic peak areas of isotopically labeled amino acids and internal standards. High-confidence metabolites were those which had expected behavior in a dilution series and a coefficient of variation below the defined threshold.

### Data analysis – Pyxis

Absolute quantitation via the Pyxis platform was performed using a web-based application that runs the Pyxis model. Raw data files were uploaded to the cloud (Amazon S3). A metadata file with their S3 locations was then uploaded to the web application, which converts the files to mzML format and runs the Pyxis model using those files. Concentration data were returned in csv format. The reported values are adjusted for the dilution factor (10x) used. For datasets in which Pyxis concentration data is compared to peak area data, peak areas were quantified using TraceFinder.

### Orthogonal concentration determination with heavy isotopologues

Matched heavy isotopologues were purchased from Cambridge Isotope Laboratories, Inc. Heavy isotopologues were added into the extraction solution at a final concentration of 5 µM for each analyte. Quantitation of peak areas for the matched heavy and light metabolites was performed with TraceFinder. Converting peak area to concentration was performed using the formula below.

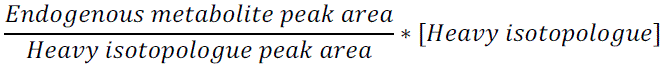

### Untargeted analysis

Metabolite identification using MS/MS data was performed in MZmine3^29^. A cosine similarity score of 0.7 was used to identify high-confidence matches. Spectral libraries from GNPS^37^ and the MassBank of North America were used in MZmine.

### Pyxis Model Architecture and Training

The Pyxis model uses a transformer-based architecture to predict analyte concentrations from mass spectrometry data. The model consists of an encoder that processes molecular features and intensity data from positive and negative ion modes. Molecular features are encoded using a linear layer, while intensity data is processed through a 1D patchification layer. These embeddings are combined with learned positional encodings and passed through a transformer encoder with 6 layers and 6 attention heads. The final hidden state is used to predict analyte concentration through a multi-layer perceptron head. The model was trained on a dataset of mass spectrometry samples with known analyte concentrations. Training utilized a weighted random sampler to balance representation across analytes and concentration ranges. Data augmentation techniques including random translation, channel dropout, and magnitude scaling were applied during training. The model was optimized using AdamW with a one-cycle learning rate schedule and gradient clipping. Training was performed for 250 epochs using mixed-precision and distributed data parallel training across 8 GPUs. Model checkpoints were saved based on validation loss and weighted mean absolute percentage error metrics.

### Statistical Analysis

All data analysis and plotting were performed in Python. For the linearity filter, fitting was performed with SciPy. Filter criteria were as follows: slope > 0.7, R-squared value > 0.5, intercept between –0.3 and 0.3.

### Data Availability

Raw mass spectrometry data generated as part of this study is deposited at Metabolomics Workbench (Study ID #ST003441).

### Code Availability

Code relating to the Pyxis model architecture is proprietary. Access to Pyxis can be obtained through a commercial agreement with Matterworks, Inc.

## Acknowledgments

We thank all members of Matterworks and the Kanarek lab for their advice and help. We are grateful for the following support: Canadian Institutes of Health Research Fellowship (A.Y.L.W.), BCH Pilot Grant (N.K.), and BCH BTREC Fund (N.K.). N.K. is a Pew Biomedical Scholar.

## Author Contributions

A.Y.L.W. performed all animal work. L.S.F., J.H., and A.Y.L.W. prepared samples and collected mass spectrometry data. L.S.F. and A.Y.L.W. performed data analysis. L.S.F., A.Y.L.W., J.M.C., and N.K. wrote the manuscript. L.S.F., J.M.C., A.S.H.C, J.G.P., J.D.L. and S.B.H. collected and curated the training dataset for Pyxis. T.K. and D.S. wrote the code for and trained the Pyxis model. M.C.D. conceived the study. All authors have read and agreed to the published version of the manuscript.

---

**Supplementary Fig. 1.**
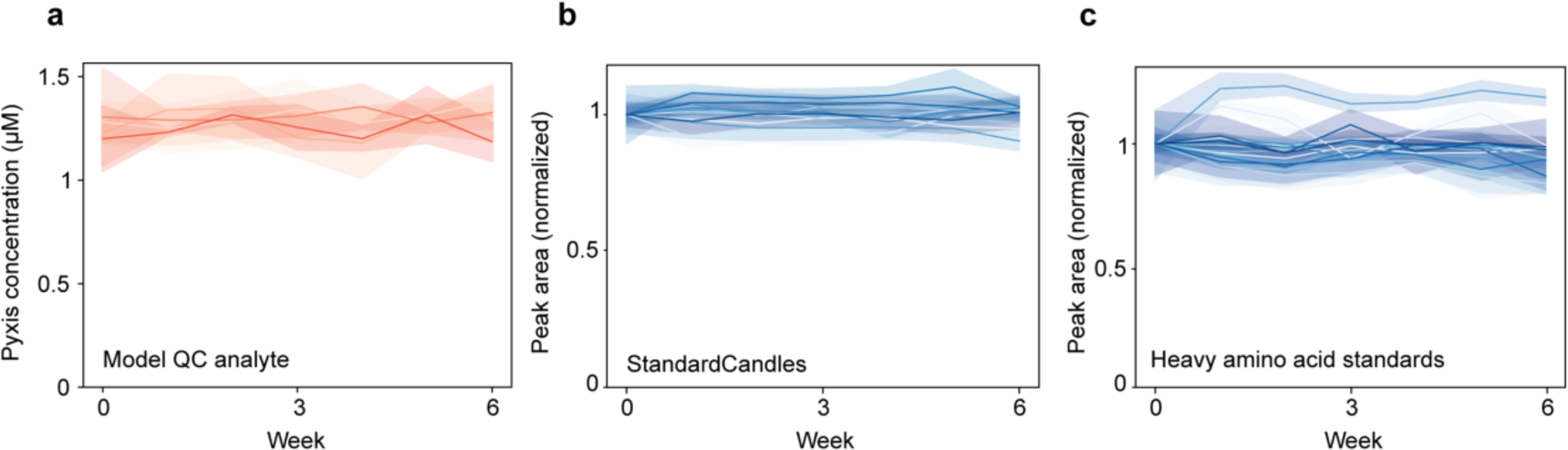
Quality control data relating to the Pyxis LC-MS method. **a** A proprietary QC analyte is spiked into the extraction buffer at 1 µM and quantified by Pyxis. Different sample pools are shown in different shades of red. **b-c** QC plots using peak areas from the StandardCandles mixture (**b**) and a set of spiked-in heavy isotopologues (**c**). Peak areas are normalized to the week 0 signal for each analyte. Different analytes are depicted with different shades of blue. The shaded area depicts standard deviation between 2 technical replicates.

**Supplementary Fig. 2.**
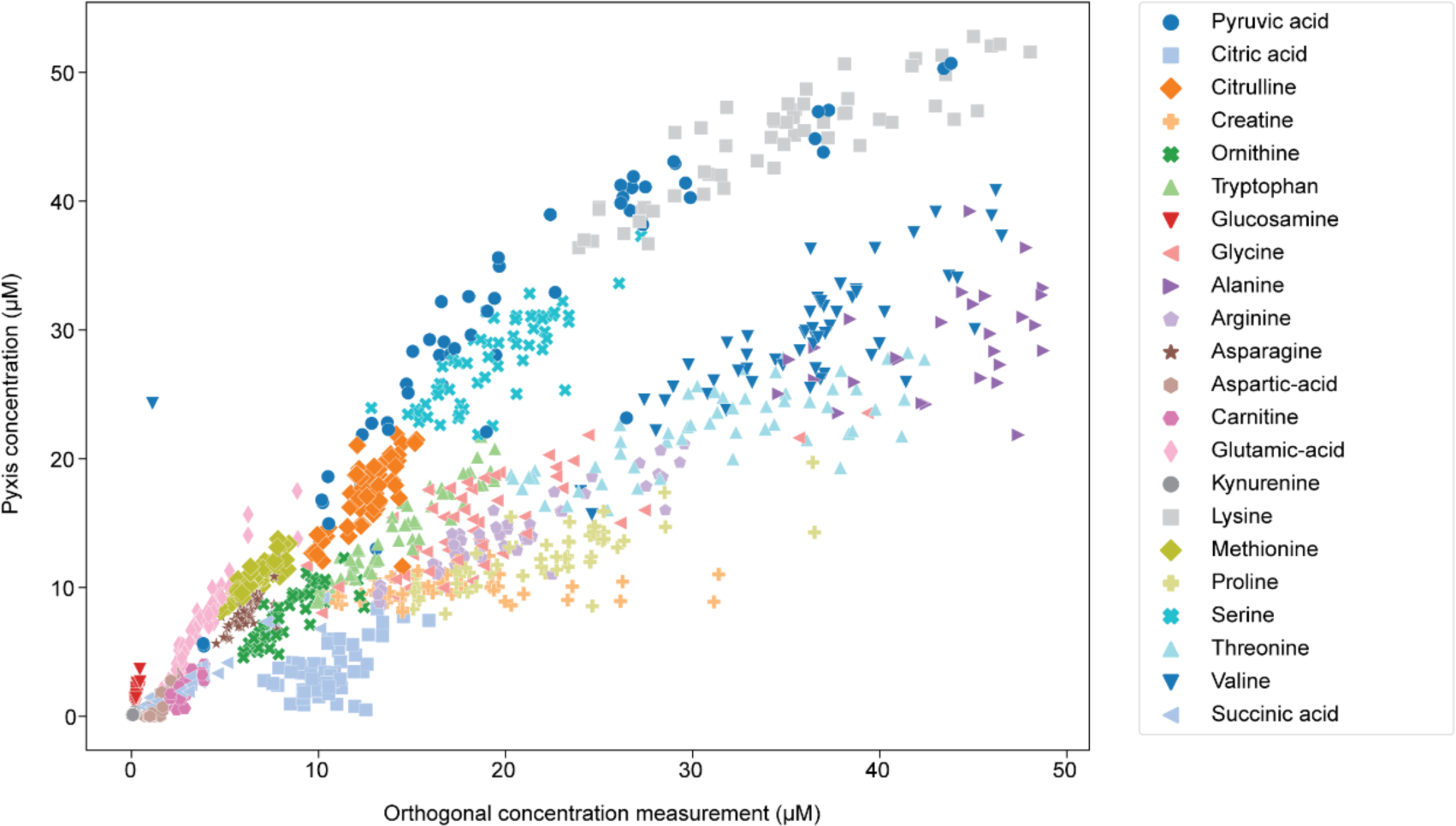
Comparison of Pyxis concentration data to orthogonal concentration measurements for 22 endogenous metabolites present in the plasma samples. Data are related to Fig. 2a.

**Supplementary Fig. 3.**
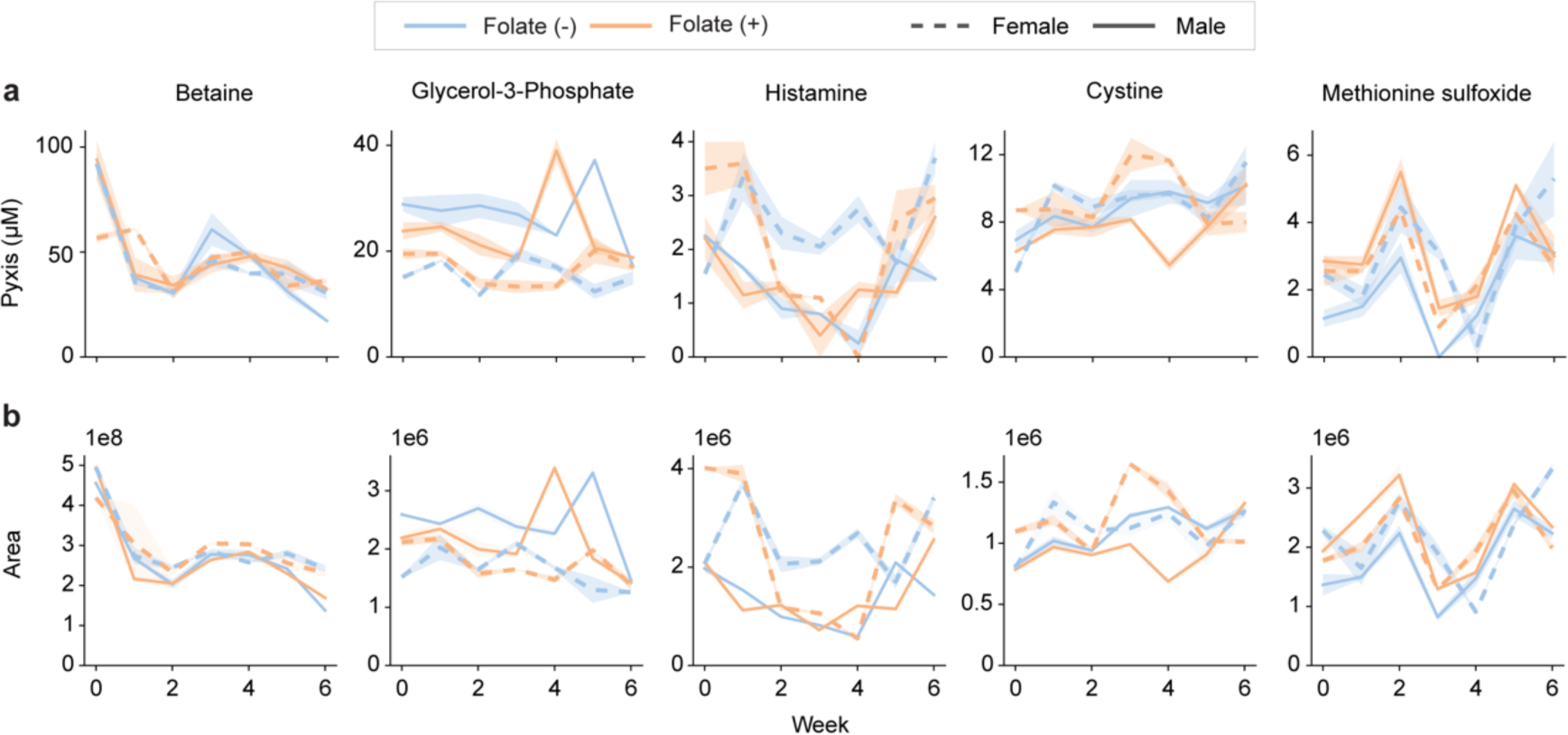
Comparison of Pyxis concentration measurements (**a**) to peak area measurements (**b**) for a set of metabolites. Legend given above the plots. The shaded area depicts standard deviation between 2 technical replicates.

**Supplementary Fig. 4.**
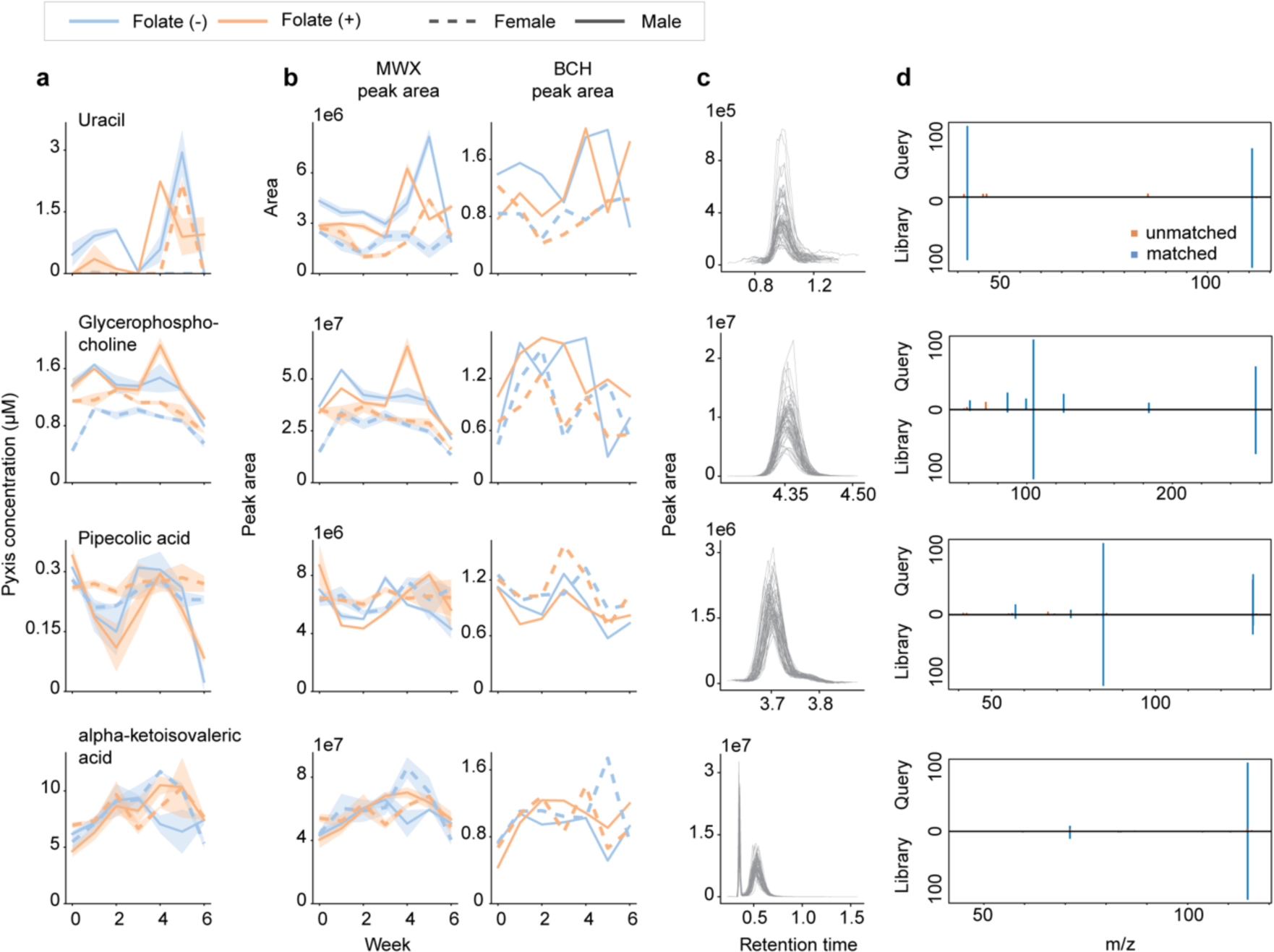
Additional examples relating to Fig. 3. **a** Pyxis concentration measurement for each metabolite. Legend given above the plots. **b** Peak area measurements (arbitrary units) given for Matterworks (MWX) and BCH. The shaded area depicts standard deviation between 2 technical replicates. **c** Base peak plots for each metabolite in the pool sample. **d** MS/MS data mirror plots for each analyte. Top half is the query (acquired) spectrum, bottom half is the library spectrum. Blue bars are those fragments in which there was a library match. Orange bars are unmatched fragments. Data were acquired from pooled plasma samples.

**Supplementary Fig. 5.**
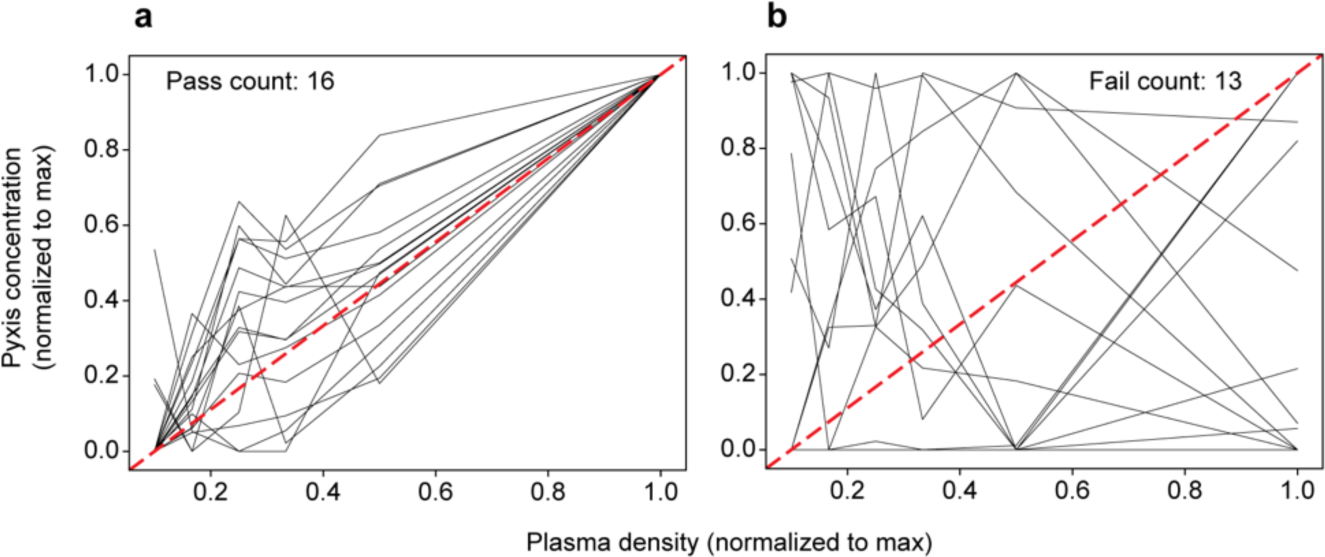
Pooled plasma sample dilution series data on the 29 metabolites that were included in the relative quantitation dataset but not the original Pyxis data and were detected by the Pyxis LC-MS method. **a** 16 metabolites passed the linearity filter (described in Methods). **b** 13 metabolites did not pass the filter. The red dotted line represents a y=x line. Each metabolite is plotted as a separate line.

**Supplementary Table 1.** Table depicting the volumes of plasma in µL collected from each mouse (top) and in each pool (bottom). Pools 1-4 consisted of samples drawn from 4, 3, 4 and 3 mice, respectively.

|  | Sex | Diet | Week 0 | Week 1 | Week 2 | Week 3 | Week 4 | Week 5 | Week 6 |
| --- | --- | --- | --- | --- | --- | --- | --- | --- | --- |
| Pool 1 | F | Folate Replete | 10 | 15 | 15 | 15 | 15 | 10 | 15 |
|  | F | Folate Replete | 15 | 5 | 15 | 15 | NA | NA | NA |
|  | F | Folate Replete | 15 | 15 | 10 | 15 | 10 | 10 | 15 |
|  | F | Folate Replete | 15 | 15 | 15 | 10 | NA | NA | NA |
| Pool 2 | F | Folate Deficient | 15 | 10 | 15 | 5 | 15 | 15 | 10 |
|  | F | Folate Deficient | 10 | 10 | 15 | 15 | NA | NA | NA |
|  | F | Folate Deficient | 15 | 15 | 15 | 15 | 5 | 15 | 10 |
| Pool 3 | M | Folate Deficient | 15 | 10 | 15 | 10 | 15 | 15 | 10 |
|  | M | Folate Deficient | 15 | 10 | 15 | 15 | 15 | 15 | 15 |
|  | M | Folate Deficient | 10 | 15 | 15 | 10 | NA | NA | NA |
|  | M | Folate Deficient | 15 | 5 | 5 | 15 | NA | NA | NA |
| Pool 4 | M | Folate Replete | 15 | 15 | 15 | 10 | 15 | 15 | 15 |
|  | M | Folate Replete | 5 | 5 | 15 | 15 | 5 | 15 | 10 |
|  | M | Folate Replete | 15 | 5 | 15 | 15 | NA | NA | NA |
| Total volume |  |  |  |  |  |  |  |  |  |
| Pool 1 | F | Folate Deficient | 55 | 50 | 55 | 55 | 25 | 20 | 30 |
| Pool 2 | F | Folate Replete | 40 | 35 | 45 | 35 | 20 | 30 | 20 |
| Pool 3 | M | Folate Deficient | 55 | 40 | 50 | 50 | 30 | 30 | 25 |
| Pool 4 | M | Folate Replete | 35 | 25 | 45 | 40 | 20 | 30 | 25 |

**Supplementary Table 2.**
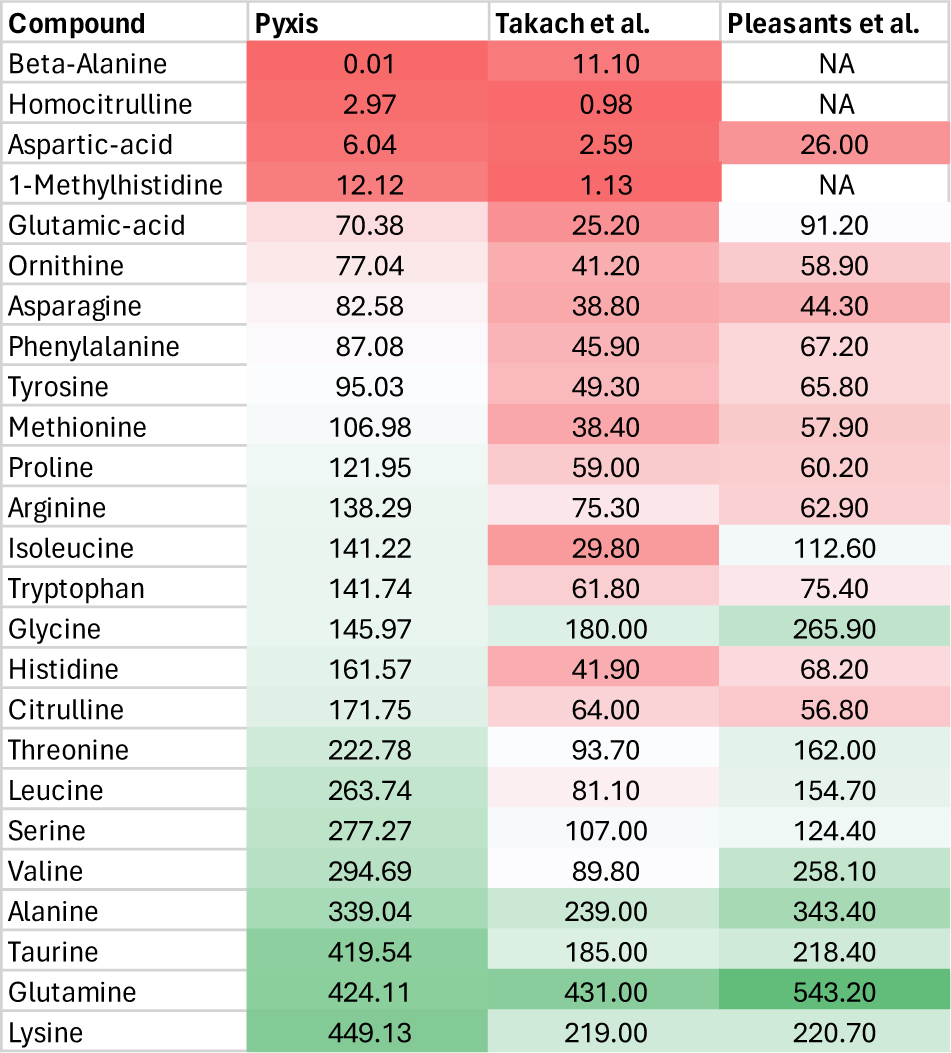
Table depicting the mouse plasma concentrations (μM) reported by Pyxis (this study), Takach et al. ^25^, and Pleasants et al. ^26^. Table is colored by concentration value where green is high concentration and red is low concentration (relative to the median).

| Compound | Pyxis | Takach et al. | Pleasants et al. |
| --- | --- | --- | --- |
| Beta-Alanine | 0.01 | 11.10 | NA |
| Homocitrulline | 2.97 | 0.98 | NA |
| Aspartic-acid | 6.04 | 2.59 | 26.00 |
| 1-Methylhistidine | 12.12 | 1.13 | NA |
| Glutamic-acid | 70.38 | 25.20 | 91.20 |
| Ornithine | 77.04 | 41.20 | 58.90 |
| Asparagine | 82.58 | 38.80 | 44.30 |
| Phenylalanine | 87.08 | 45.90 | 67.20 |
| Tyrosine | 95.03 | 49.30 | 65.80 |
| Methionine | 106.98 | 38.40 | 57.90 |
| Proline | 121.95 | 59.00 | 60.20 |
| Arginine | 138.29 | 75.30 | 62.90 |
| Isoleucine | 141.22 | 29.80 | 112.60 |
| Tryptophan | 141.74 | 61.80 | 75.40 |
| Glycine | 145.97 | 180.00 | 265.90 |
| Histidine | 161.57 | 41.90 | 68.20 |
| Citrulline | 171.75 | 64.00 | 56.80 |
| Threonine | 222.78 | 93.70 | 162.00 |
| Leucine | 263.74 | 81.10 | 154.70 |
| Serine | 277.27 | 107.00 | 124.40 |
| Valine | 294.69 | 89.80 | 258.10 |
| Alanine | 339.04 | 239.00 | 343.40 |
| Taurine | 419.54 | 185.00 | 218.40 |
| Glutamine | 424.11 | 431.00 | 543.20 |
| Lysine | 449.13 | 219.00 | 220.70 |

